# *In vivo* silencing of regulatory elements using a single AAV-CRISPRi vector

**DOI:** 10.1101/2023.11.05.565512

**Authors:** P Laurette, C Cao, D Ramanujam, M Schwaderer, T Lueneburg, S Kuss, L Weiss, R Dilshat, EEM Furlong, F Rezende, S Engelhardt, R Gilsbach

**Affiliations:** Institute of Experimental Cardiology, Heidelberg University Hospital, Heidelberg, Germany; DZHK (German Center of Cardiovascular Research) Partner Site Rhein/Main, München and Heidelberg/Mannheim, Germany; Institute of Cardiovascular Physiology, Frankfurt University, Frankfurt, Germany; Institute of Pharmacology and Toxicology, Technical University of Munich, Munich, Germany; Institute of clinical and experimental Pharmacology and Toxicology, University of Freiburg, Germany; European Molecular Biology Laboratory (EMBL), Genome Biology Unit, Heidelberg, Germany

## Abstract

CRISPR-Cas9 based transcriptional repressors (CRISPRi) have emerged as specific and robust tools for functional epigenetic silencing of regulatory elements. Adeno-associated viruses (AAVs) are promising CRISPRi delivery vectors for cardiovascular research and therapy. However, compact vectors enabling codelivery of all CRISPRi components by a single AAV are needed for an enhanced and consistent performance.

We engineered a 4.7kb all-in-one CRISPRi construct compatible with AAV-mediated delivery and produced cardiotropic AAVi 6 and 9 particles for in *vitro* and in *vivo* application, respectively.

AAVi vectors targeting the *Nppa* promoter (AAVi^Nppa^) reduced gene expression in cultivated cardiomyocytes (HL-1 cells) in a dose-dependent manner. The maximum effect was a >95% reduction as measured by qPCR and RNA-seq. This effect was orchestrated by loss of chromatin accessibility (ATAC-seq) and establishment of heterochromatin (H3K9me3 ChIP-seq) specifically at the target promoter region. We confirmed the broad applicability of AAVi to different cardiomyocyte cell culture systems by silencing several genes in primary neonatal rat ventricular cardiomyocytes (NRVMs), human iPSC-derived cardioids and iPSC-CMs.

To demonstrate the efficacy of AAVi in *vivo* we injected 8-week-old C57Bl/6 WT mice with a single dose of AAVi^Nppa^ and implanted osmotic minipumps releasing Phenylephrine (50 mg/kg/d) and Angiotensin II (0.45 mg/kg/d) to induce *Nppa* transcription. AAVi^Nppa^ silenced *Nppa* transcription as revealed by qPCR and single nuclei RNA-seq even under stress conditions. On the epigenome layer AAVi^Nppa^ induced closed chromatin at the *Nppa* promoter site comparable to the in *vitro* effect. Here, we present an efficient AAV-based method for CRISPRi-mediated epigenetic silencing of gene expression in cardiac myocytes *in vivo* and *in vitro*. This functional epigenetic approach provides an efficient way to modulate gene expression in the heart and could become a standard method for cardiovascular disease modelling and translational research.

## Introduction

CRISPR-Cas9 based transcriptional repressors (CRISPRi) have emerged as powerful tools for functional epigenetic silencing ^1^. They rely on a nuclease-deficient Cas9 fused to a repressor domain (dCas9i) and a target-specific guide RNA (gRNA). In contrast to gene editing approaches, dCas9-KRAB mediated CRISPRi is generally reversible, titratable and circumvents the risks associated with direct modification of the genomic sequence.

Adeno-associated viruses (AAVs) are promising non-integrating vectors for gene therapy, but their limited packaging capacity make them incompatible with large inserts such as the common *S. pyogenes* dCas9i. Here we describe a CRISPR-based gene silencing method in the heart relying on a single compact vector compatible with adeno-associated virus delivery (AAVi).

## Results and Discussion

We engineered an AAV vector containing both a *S. aureus*-derived nuclease-deficient Cas9 fused to a KRAB repressor domain ^2^ driven by a CMV promoter and a gRNA cassette under the control of the U6 promoter (Fig. 1A). The CMV promoter ensures robust expression and can be exchanged if cell type-specific expression is required. Organ and cell type selectivity can further be modulated by the selected capsid serotypes as shown for muscle tissue ^3^. The insert size of the engineered AAVi construct is 4.7kb reaching the upper limit insert size of the AAV genome. Nevertheless, the yields of AAV6 and AAV9 particles were comparable to those produced using an AAV vector with smaller (3kb) insert size (Fig. 1B).

**Figure 1:**
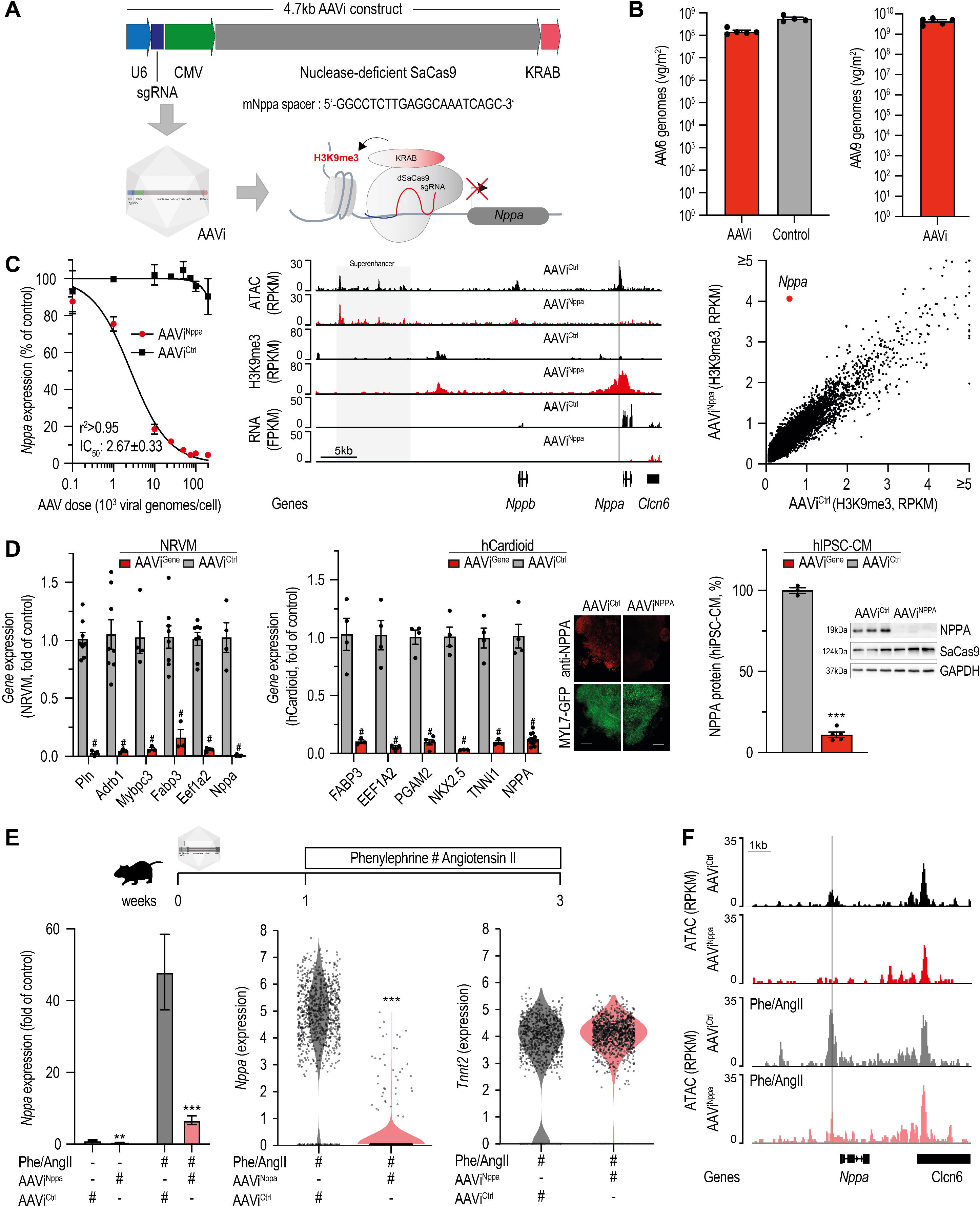
Functional epigenetic promoter silencing using a single AAV-CRISPRi vector. (A) Schematic representation of the CRISPRi construct, generation of AAVi particles and CRISPRi mediated silencing. The spacer sequence used for silencing of *Nppa* in mouse cardiomyocytes (CM) is indicated. (B) AAV6 or 9 particles were generated for the CRISPRi and reference AAV plasmids (insert size 3kb) and the resulting viral genomes were quantified by RT-qPCR of AAV-ITRs (bar chart, n=3-5). (C) Increasing doses of AAV6 particles containing a non-targeting construct (AAVi^Ctrl^) or one targeting the *Nppa* promoter (AAVi^Nppa^) were used to transduce HL-1 mouse cardiac myocytes *in vitro* and gene expression was measured by RT-qPCR. A non-linear regression was used to determine the half-maximal effect (IC_50_ ± SEM) and the goodness of fit (r^2^) (left plot, n=3). The effect of AAVi^Nppa^ on chromatin accessibility (ATAC-seq), heterochromatin formation (H3K9me3) and gene expression (RNA-seq) in HL-1 cells is displayed for the *Nppa* locus (middle plot). A genome-wide analysis of H3K9me3 deposition in AAVi^Nppa^ vs. AAVi^Ctrl^ is shown for all annotated gene promoters (+/-2.5kb) (right plot). (D) AAVi was used to silence multiple genes in neonatal rat cardiomyocytes NRVM (left panel), human cardioids (middle panel) as well as human iPSC-CMs (right panel). A single spacer sequence was used for each gene except both human and rat NPPA for which we designed two and merged data. Gene expression was assessed by RT-qPCR (n=3-12). Immunohistochemistry was used to assess NPPA protein levels (red signal) in human cardioids. Cardiomyocytes were visualized using GFP-tagged MYL7 (green signal). NPPA protein levels were quantified using western blot analysis in human iPSC-CM (n=3-5). Representative western blot data are shown for NPPA, SaCas9 and GAPDH (loading control). (E) Timeline of AAVi *in vivo* silencing experiments. Phenylephrine- and Angiotensin II-releasing osmotic mini pumps were used to induce cardiac hypertrophy. RT-qPCR analysis of *Nppa* gene expression in ventricular tissue (left panel, n=7). Single nuclear RNA-seq analysis of *Nppa* and *Tnnt2* expression. Data shown were obtained from 1165 (AAVi^Nppa^) and 913 (AAVi^Ctrl^) cardiomyocyte nuclei. (F) Original traces of chromatin accessibility (ATAC-seq) in sorted cardiomyocyte nuclei. Shown are mean ± SEM. *** p<0.001, ** p<0.01, ANOVA; ^#^ q<0.01, FDR t-Test, Statistical analysis of ratios was performed after log-transformation for normal distribution.

To show the functionality of AAVi for cardiac research we silenced the natriuretic peptide A (NPPA) since NPPA is highly and specifically expressed in cardiomyocytes (CM). We designed gRNAs targeting the accessible *Nppa* proximal promoter region (AAVi^Nppa^, Fig.1A) or without homology to mammalian genomes (AAVi^Ctrl^). We packaged them into AAV6 particles to transduce cultured mouse HL-1 cardiomyocytes. AAVi^Nppa^ reduced *Nppa* gene expression in HL-1 cells in a dose-dependent manner after one week (Fig. 1C). Due to cell proliferation associated AAV dilution the effect is then progressively lost (data not shown) as AAVi mediated silencing is not heritable. Epigenome analysis using ATAC-seq and ChIP-seq showed local loss of chromatin accessibility and a stretch of H3K9me3 ±2kb around the target site, orchestrating loss of transcriptional activity (RNA-seq) and confirming successful epigenetic silencing (Fig. 1C). A genome wide analysis of gene promoters revealed that *de novo* deposition of the heterochromatin mark was specific to the target promoter region of *Nppa* and did not spread to known interacting regions such as the *Nppb* promoter and the shared superenhancer ^4^ (Fig. 1C). One H3K9me3 positive off-target site was identified at a non-regulatory site of the genome homologous with the 3’half of the spacer. We then confirmed the broad applicability of AAVi to different cardiomyocyte cell culture systems by silencing several genes in primary neonatal rat ventricular cardiomyocytes (NRVMs), human iPSC-derived cardioids and iPSC-CMs (Fig.1D). For human iPSC-derived cells we confirmed the effect of AAVi on NPPA RNA and protein levels, as representatively shown for cardioids by immunofluorescence and quantitatively for iPSC-CM by western blot analysis. In total 80% (17 out of 21) of tested sgRNAs permitted efficient gene silencing documenting the high efficacy of AAVi.

*Nppa* transcription is induced during cardiac stress and heart disease. To demonstrate the robustness of AAVi *in vivo*, we aimed to antagonize this process in an experimental heart disease model. Therefore, we retro-orbitally injected 8-week-old male C57Bl/6 mice with a single dose of AAVi^Nppa^ or AAVi^Ctrl^ (n=7, 3x10E12 AAV9 VG/mouse). 7 days later we implanted osmotic minipumps releasing Phenylephrine (50 mg/kg/d) and Angiotensin II (450 μg/kg/d) over two weeks to mimic neurohormonal stress signals and induce cardiac hypertrophy. AAVi^Nppa^ resulted in >7-fold downregulation of ventricular *Nppa* expression (Fig. 1E). Single nuclei RNA-seq analysis of cardiac nuclei further revealed that *Nppa* expression is detectable by snRNA-seq in 3.5% (41/1165) of cardiomyocyte nuclei (1151/1165 Tnnt2^+^) from mice treated with AAVi^Nppa^ as compared to 95% (864/913) upon AAVi^Ctrl^ (770/913 Tnnt2^+^), demonstrating homogenous downregulation of *Nppa* (Fig. 1E). Notably, the efficacy of AAVi exceeded a previously study reporting a 32% reduction of *Pcsk9* mRNA expression in mouse liver using an alternative single AAV-CRISPRi system ^5^. To show the effect on the epigenetic layer and the specificity of AAVi, we sorted cardiomyocyte nuclei from diseased hearts and analyzed changes in chromatin accessibility (ATAC-seq) and gene expression (nuclear RNA-seq) on a genome-wide scale. AAVi^Nppa^ established closed chromatin at the *Nppa* promoter (Fig. 1F) and *Nppa* was downregulated, while accessibility and expression of neighboring genes weren’t affected. Beside Nppa, we found only 25 genes mainly linked to antiviral response to be affected (n=3, q ≤ 0,001).

## Summary

Here, we present an efficient AAV-based method for CRISPRi-mediated epigenetic silencing of gene expression in cardiac myocytes *in vivo* and *in vitro*. This functional epigenetic approach provides a novel and efficient way to modulate gene expression in the heart and could become a standard method for cardiovascular disease modelling and translational research.

## Acknowledgments

We thank Sabine Brummer, Joshua Hartmann and Núria Diaz i Pedrosa for their technical assistance and Benjamin Meder (Institute for Cardiomyopathies, Heidelberg) for providing access to Novaseq6000. We thank Sasha Mendjan (IMBA, Vienna, Austria and Mercator fellow of the CRC1550) for help with establishing the generation of cardioids in the lab of EEMF. The authors acknowledge the support of the Freiburg Galaxy Team: Rolf Backofen and Björn Grüning, Bioinformatics, University of Freiburg (Germany) funded by the German Federal Ministry of Education and Research (grant 031 A538A de.NBI-RBC) and the CRC1425 (S03). CC was supported by the China Scholarship Council. This study was supported by the Collaborative Research Centers 1425 (projects P02 and S03 to RG) and 1550 (project A02 to RG, A04 to EEMF), the Rolf M. Schwiete Stiftung (Mannheim, Germany) to RG and the DZHK (project 81X450019 to SE, DR, PL, RG). The authors acknowledge the data storage service SDS@hd supported by the Ministry of Science, Research and the Arts Baden-Württemberg (MWK) and the German Research Foundation (DFG) through grant INST 35/1503-1 FUGG. SE is supported by the DFG (TRR267) and the BMBF in the framework of the Cluster for Nucleic Acid Therapeutics (CNATM).

## References

1. Qi LS, Larson MH, Gilbert LA, Doudna JA, Weissman JS, Arkin AP, Lim WA. Repurposing CRISPR as an RNA-guided platform for sequence-specific control of gene expression. Cell. 2013;152:1173–1183. doi: 10.1016/j.cell.2013.02.022

2. Thakore PI, Kwon JB, Nelson CE, Rouse DC, Gemberling MP, Oliver ML, Gersbach CA. RNA-guided transcriptional silencing in vivo with S. aureus CRISPR-Cas9 repressors. Nat Commun. 2018;9:1674. doi: 10.1038/s41467-018-04048-4

3. Weinmann J, Weis S, Sippel J, Tulalamba W, Remes A, El Andari J, Herrmann AK, Pham QH, Borowski C, Hille S, et al. Identification of a myotropic AAV by massively parallel in vivo evaluation of barcoded capsid variants. Nat Commun. 2020;11:5432. doi: 10.1038/s41467-020-19230-w

4. Man JCK, van Duijvenboden K, Krijger PHL, Hooijkaas IB, van der Made I, de Gier-de Vries C, Wakker V, Creemers EE, de Laat W, Boukens BJ, et al. Genetic Dissection of a Super Enhancer Controlling the Nppa-Nppb Cluster in the Heart. Circ Res. 2021;128:115–129. doi: 10.1161/CIRCRESAHA.120.317045

5. Backstrom JR, Sheng J, Wang MC, Bernardo-Colon A, Rex TS. Optimization of S. aureus dCas9 and CRISPRi Elements for a Single Adeno-Associated Virus that Targets an Endogenous Gene. Mol Ther Methods Clin Dev. 2020;19:139–148. doi: 10.1016/j.omtm.2020.09.001

